# Deterministic shifts in molecular evolution correlate with convergence to annualism in killifishes

**DOI:** 10.1101/2021.08.09.455723

**Authors:** Andrew W. Thompson, Amanda C. Black, Yu Huang, Qiong Shi, Andrew I. Furness, Ingo Braasch, Federico G. Hoffmann, Guillermo Ortí

**Affiliations:** Department of Integrative Biology, Michigan State University, East Lansing, Michigan 48823, USA; Ecology, Evolution & Behavior Program, Michigan State University, East Lansing, MI, USA; Department of Biochemistry, Molecular Biology, Entomology, & Plant Pathology, Mississippi State University, Starkville, MS 39759, USA; Shenzhen Key Lab of Marine Genomics, Guangdong Provincial Key Lab of Molecular Breeding in Marine Economic Animals, BGI Marine, Shenzhen 518083, China; BGI Education Center, University of Chinese Academy of Sciences, Shenzhen 518083, China; Department of Biological Sciences, The George Washington University, Washington, DC 20052, USA; Department of Biological and Marine Sciences, University of Hull, UK

**Keywords:** adaptation to extreme environments, determinism, diapause, OXPHOS, phylogenetics

## Abstract

The repeated evolution of novel life histories correlating with ecological variables offer opportunities to test scenarios of convergence and determinism in genetic, developmental, and metabolic features. Here we leverage the diversity of aplocheiloid killifishes, a clade of teleost fishes that contains over 750 species on three continents. Nearly half of these are “annual” or seasonal species that inhabit bodies of water that desiccate and are unfeasible for growth, reproduction, or survival for weeks to months at a time. We present a large-scale phylogenomic reconstruction of aplocheiloid killifish evolution using newly sequenced transcriptomes from all major clades. We show that developmental dormancy (diapause) and annualism have up to seven independent origins in Africa and America. We then measure evolutionary rates of orthologous genes and show that annual life history is correlated with higher d*N*/d*S* ratios. Many of these fast-evolving genes in annual species constitute key developmental genes and nuclear-encoded metabolic genes that control oxidative phosphorylation. Lastly, we compare these fast-evolving genes to genes associated with developmental dormancy and metabolic shifts in killifishes and other vertebrates and thereby identify molecular evolutionary signatures of repeated evolutionary transitions to extreme environments.

## Introduction

Convergent and parallel evolution are active and important focal areas of research in evolutionary biology. Multiple independent origins or losses of phenotypes act as evolutionary replicates to investigate the genomic bases of diverse cases of morphological, physiological, and/or behavioral adaptations associated with environmental challenges and transitions. One outstanding question is whether or not evolution is deterministic and to what degree similar traits evolve predictably and repeatedly across similar environment conditions (Gould 1989; Losos 1998). This question can be addressed by studying radiations of both distantly and closely related organisms across common environmental gradients (Chen et al. 1997; Greenway et al. 2020; Rincon-Sandoval et al. 2020). Such traits can manifest at the molecular, morphological, developmental, and/or physiological levels. Examination of their multiple origins and losses can give insight into the evolution of novelty, evolvability, and predictability of adaptive optima. In the genomics era, we can now explore the functional genomics of key traits with increased power to resolve phylogenies with confidence.

To test whether life history repeatedly correlates with changes in rates of molecular evolution, we leverage the diversity of aplocheiloid killifishes that include lineages thought to have convergently evolved multiple times the ability to live in extreme, temporary environments (Parenti 1981; Murphy and Collier 1997; Costa 2013; Furness et al. 2015). Aplocheiloid killifishes represent a diverse group of vertebrates with over 780 valid species (Fricke et al. 2021), roughly half of which live in ephemeral habitats with extreme seasonal variation that desiccate once or twice per year, or intermittently, in tropical regions of Africa and South America (Costa 2002; Furness 2015). Thus, depending on the species and climate, killifishes can complete one (annual) or two (bi-annual) life cycles per year, hence the term “annualism”, (henceforth used to describe seasonality). Desiccation results in the death of the adult population (Myers 1942; Simpson 1979), but the next generation survives as dormant, diapausing embryos (Wourms 1972a; Wourms 1972b; Wourms 1972c) within specialized eggs (Thompson et al. 2017a) that are resistant to extreme environmental conditions. Often, annual species are subject to rapid senescence due to relaxation of selection on longevity, making them increasingly popular biomedical model species for aging research (Jagadeeswaran 2006; Harel, Benayoun, B.E. Machado, et al. 2015; Wendler et al. 2015; Hu and Brunet 2018; Cui et al. 2019; Reuter et al. 2019). Additionally, seasonal killifishes and their non-seasonal relatives represent a “model clade” (Sanger and Rajakumar 2019; Jourjine and Hoekstra 2021) to explore the evolution of life in extreme environments. Much less is known about the molecular mechanisms that killifish use to control dormancy, desiccation tolerance, and how these developmental and physiological adaptations evolved in more than one family of killifishes.

Annual killifish embryos undergo up to three distinct diapause stages (DI, DII, DIII) during development (Wourms 1972a; Wourms 1972b; Wourms 1972c) that can last for months to years (Hu et al. 2020). Diapause I occurs during blastomere development, DII during somitogenesis, and DIII prior to hatching after organogenesis is completed (Wourms 1972a; Wourms 1972b; Wourms 1972c). Diapause I illustrates a completely novel pattern of cell migration where blastomeres become ameboid and move around before their coalescence and the resumption of development (Carter and Wourms 1990; Arezo et al. 2017; Dolfi et al. 2019). Diapause II confers the most extreme resistance to the embryos for many physiological stressors such as desiccation (Podrabsky et al. 2001; Podrabsky et al. 2010; Daniel et al. 2020), DNA damage (Wagner and Podrabsky 2014) and anoxia (Podrabsky et al. 2007; Podrabsky and Wilson 2016; Riggs et al. 2019). Diapause II is the defining feature of killifish “annualism”. Diapause II can be escaped via changes in temperature and vitamin D signaling (Romney et al. 2018), resulting in direct development to DIII. Diapause III allows the embryo to control hatching and is associated with the expression of genes shown to be convergently expressed during dormancy across several metazoan lineages (Thompson and Ortí 2016). Diapause III is the only vertebrate diapause known to occur after organogenesis allowing fully formed individuals to enter a state of suspended animation that has a different transcriptional profile from that of delayed-hatching embryos in related killifishes (Thompson et al. 2017b). When the habitat is flooded, embryos escape DIII and hatch (Wourms 1972a; Wourms 1972b; Wourms 1972c). Upon hatching, however, annual killifish larvae and juveniles can develop rapidly to become sexually mature in a matter of weeks (Vrtílek et al. 2018).

Until now, all molecular evolutionary studies on killifish diapause or aging either focus on a single species (Reichwald et al. 2015; Valenzano et al. 2015; Thompson and Ortí 2016; Wagner et al. 2018) or a subclade of killifishes (Cui et al. 2019) or take advantage of only a few genetic loci (Murphy and Collier 1997; Hrbek and Larson 1999; Furness et al. 2015; Helmsetter et al. 2016). Therefore, much of what is known about diapause in killifishes is restricted to detailed studies of a few, phylogenetically dispersed killifish model species such as *Austrofundulus limnaeus, Austrolebias sp*., *Nematolebias whitei*, and *Nothobranchius sp*. (Thompson and Ortí 2016; Arezo et al. 2017; Romney et al. 2018; Riggs et al. 2019; Hu et al. 2020; Zajic et al. 2020; Chalar et al. 2021; Polačik et al. 2021). Thus, little is known about most killifish species exhibiting the annual life cycle and a broad comparative study with an in-depth genome-wide exploration remains missing.

It has been hypothesized that diapause and annualism have originated or been lost multiple times among killifishes as the result of convergent or parallel evolution (Parenti 1981; Murphy and Collier 1997; Costa 2013; Furness et al. 2015) but this hypothesis has never been tested based on an aplocheiloid phylogeny generated from genome-wide sequencing of hundreds of loci. Questions further remain about the monophyly of annualism, specifically in South American killifishes (Furness et al. 2015). While insightful, previous findings show poorly supported or unstable relationships regarding the monophyly of new world non-annual species. It has been suggested that New World killifishes have a single origin of annualism, and that non-annual lineages are monophyletic (Murphy and Collier 1996; Huber 2012). Consequently, inferring a robust phylogeny encompassing all major killifish lineages is a critical prerequisite for investigating the evolutionary and mechanistic basis of life history adaptation to extreme environmental change observed among killifishes.

Given their unique biology, we hypothesize that annual killifish lineages have higher rates of molecular evolution compared to non-annual killifishes and that these changes would reflect determinism and repeated evolution of the same adaptive optima. More specifically, we hypothesized that annual killifish reflect this convergence with faster evolution of orthologous protein coding genes that are critical for survival in ephemeral pools. Comparative studies have revealed differences in molecular evolutionary rates among evolutionary lineages of protein families (Luz and Vingron 2006; De La Torre et al. 2015) and organisms (Lanfear et al. 2010; Saclier et al. 2018). For example, within ray-finned fishes, slow evolutionary rates have been identified in “living fossil” species such as gar, bowfin, and sturgeon (Braasch et al. 2016; Du et al. 2020; Thompson et al. 2021), while other species with numerous highly derived phenotypes like seahorses have some of the highest substitution rates among vertebrates (Lin et al. 2016).

Several hypotheses have been set forth to explain variation in substitution rates across taxa. Many correlates and causes of shifts in mutation rate have been proposed including changes in body size (Martin and Palumbi 1993), generation time (Laird et al. 1969; Bromham et al. 1996; Thomas et al. 2010), metabolic rate (Martin and Palumbi 1993; Gillooly et al. 2005), stress (Radman 1999), increased presence of environmental mutagens or radiation (Davies et al. 2004), higher temperatures (Garcia-Porta et al. 2019), efficiency of DNA repair mechanisms (Bromham 2009), mutations in polymerase genes (Cui et al. 2019), and effective population sizes (Ohta and Kimura 1971). Generally, current literature suggests that mutation rates and fixation of mutations are positively correlated with higher metabolic rate, shorter generation times, smaller population sizes, higher stress levels, higher concentrations of mutagens or radiation, and lower efficiency of DNA repair. Annual killifish species exhibit many, if not all these biological characteristics hypothesized to increase mutation rate as well as intense selection pressures that could increase the rate of protein evolution. For example, killifishes are often small-bodied (Costa 1998), have short generation times constrained by their temporary habitats (Simpson 1979), and have small, isolated populations that undergo and tolerate extreme environmental stressors (Podrabsky et al. 2007; Wagner and Podrabsky 2014; Daniel et al. 2020) in tropical climates. They also have high metabolic rates upon hatching since they grow rapidly to reach maturity and spawn before desiccation (Vrtílek et al. 2018). Thus, identifying changes in the rate of molecular evolution among killifishes may reveal genes involved in adaptation to extreme environments at multiple levels of biological organization.

Here, we take the first “omics” approach to study aplocheiloid killifish diversity worldwide, generating a phylo-transcriptomic dataset containing over 1,300 orthologous gene loci and sampling all major killifish clades across three families and three different continents. We generate a robust, time-calibrated phylogeny of the aplocheiloid clade, infer the number of origins of annualism, and explore changes in rates of protein evolution across annual lineages. We confirm multiple life history transitions with genomic scale data. We test our hypothesis that annual lineages show higher rates of protein evolution compared to non-annual fishes. We detect shifts in the ratios of nonsynonomous/synonomous (d*N*/d*S*) changes in protein coding loci to identify rapidly evolving killifish genes, showing that annual life histories are correlated with deterministic changes in protein evolution. We show that many of the genes evolving rapidly in annuals are candidates for targets of positive selection and are related to oxidative phosphorylation (OXPHOS) and developmental phenotypes, both of which need modifications to allow survival in harsh conditions. Lastly, we compare our findings to those for other vertebrate lineages to identify loci under evolution in extreme environments across clades and continents.

## Results

### Time Calibrated Killifish Phylogeny and the Evolution of Annualism

We sequenced and assembled novel reference transcriptomes from whole fish, heads, trunks, and/or whole embryos for 29 fish species (Supplementary Table 1), including 28 Cyprinodontiform species encompassing 26 Aplocheiloid killifish species and one more distantly related Atheriniform outgroup species. Thereby, we sequence representatives from most known annual and non-annual lineages of killifishes.

**Table 1.** Number of ortholog alignments with significantly different ω between annual and non-annual lineages. The number of annual lineages represented in an ortholog alignment was determined based on the most parsimonious ancestral state reconstruction. For example, if 2 annual species are present in an ortholog alignment and represent more than 2 different origins of annualism, the alignment is classified as having 2+ annual lineages.

| Group of alignments | Total # alignments | # Alignments with $\omega$ greater in annual lineages | # Alignments with significant $\omega$ difference between annual and non-annual lineages ($P < 0.05$) | # Alignments with significantly higher $\omega$ in annual lineages |
| --- | --- | --- | --- | --- |
| alignments with 1+ annual species | 1234 | 705 | 211 | 161 |
| alignments with 2+ annual lineages | 997 | 559 | 168 | 128 |
| alignments with 1 annual lineage | 237 | 146 | 43 | 33 |

Using these 29 reference transcriptomes, as well as other, previously sequenced fish genomes and transcriptomes (Supplementary Table 1) we identified 1,302 single-copy, orthologous gene loci (Supplementary Table 2). Alignments totaling 852,927 bp were used to infer a new phylogenomic hypothesis (Figure 1) with Bayesian, maximum likelihood (ML), and species tree methods. We time-calibrated our phylogeny using three divergence estimates of major fish clades. Annual lineages are highlighted in red in Figure 1. Our tree topology is highly supported, with all branch posterior probabilities of 0.99-1.0 in the Bayesian tree and largely congruent with those of other studies (Furness et al. 2015; Helmsetter et al. 2016). All three of our applied methodologies converge on the exact same topology for aplocheiloid killifishes.

**Figure 1.**
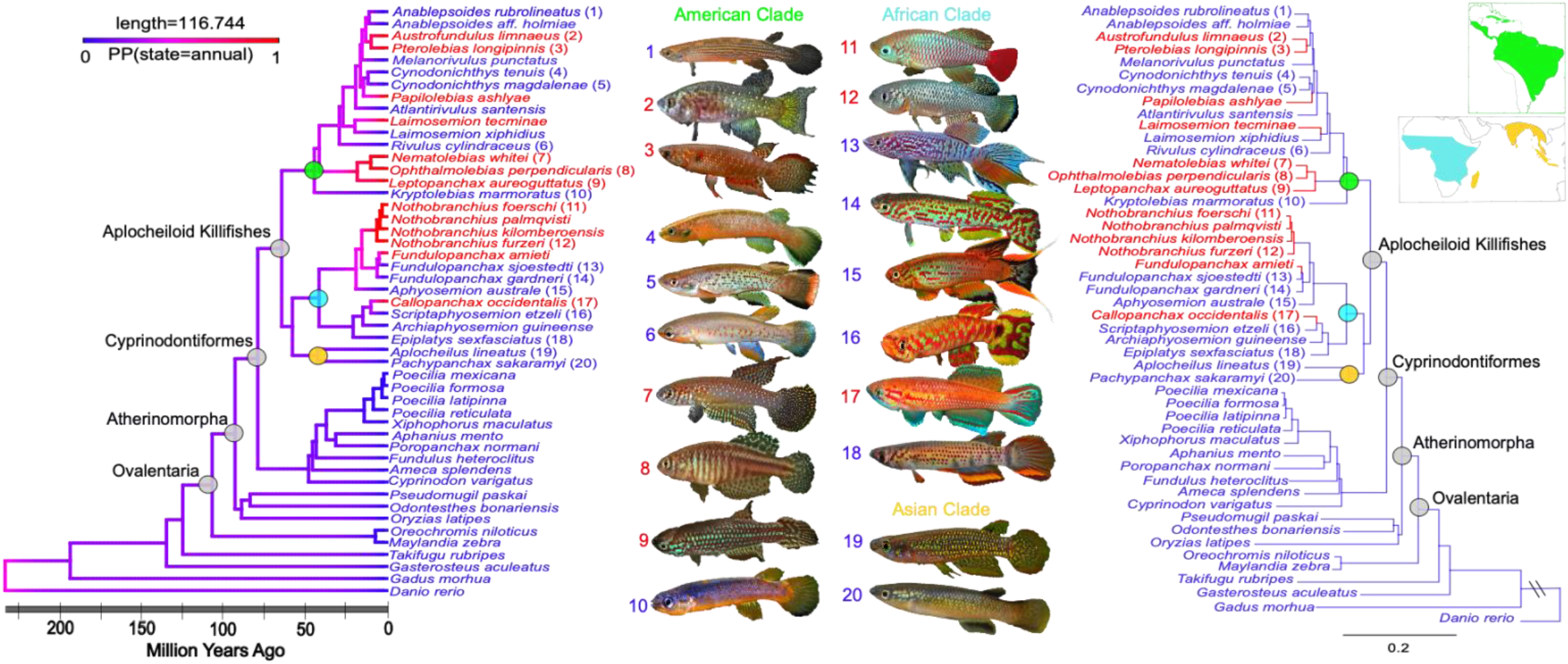
Phylogeny of Aplocheiloid killifishes. **Left:** Time calibrated, BEAST2 (Bouckaert et al. 2019) phylogeny from concatenation of 1,302 orthologous loci and 852,927 bp partitioned by codon. All support values are 0.99-1.0 posterior probability. Annual species are labeled in red. Branches are colored based on ancestral state reconstruction of annualism as a binary character inferred by an ARD model in simmap (Revell 2012) using 100,000 iterations of stochastic character mapping, and branch colors represent the posterior probability of being annual with blue as most likely to be non-annual and red most likely to be annual. Aplocheiloid killifishes began to diversify ∼64 mya. **Right:** RAxML (Stamatakis 2014) phylogeny based on concatenation of 1,302 orthologous loci and 852,927 bp partitioned by codon. Annual lineages are highlighted in red and serve as the foreground lineages for d*N*/d*S* analysis. Non-annual or background lineages are blue. Map and clade colors represent families and their geographic distribution (green = Rivulidae, Blue = Nothobranchiidae, Yellow = Aplocheilidae). All branches of aplocheiloids except the most recent common ancestor (MRCA) of *Nematolebias* and *Austrofundulus* have 100% bootstrap support (BS), and all others but *Ameca* and *Cyprinodon*; and *Fundulus* and *Poecilia* have 75%+ BS. Red clades represent the most parsimonious ancestral state reconstruction for the presence of annualism. Photos by Anthony Terceira.

**Figure 2.**
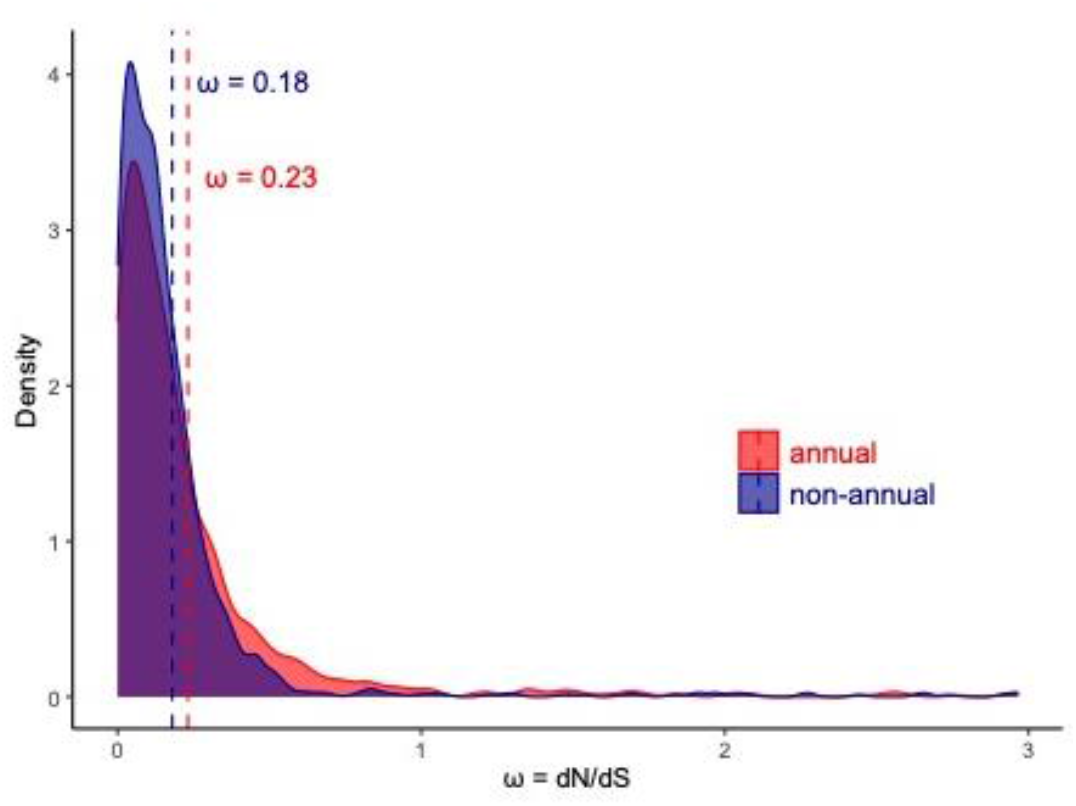
Density distribution of ω estimates for annual and non-annual species with values higher than 3 excluded. Differences between the 2 distributions were tested with Kolgomorov-Smirnoff test (D=0.10349, P = 4.515e-06).

Notably, our inference places *Kryptolebias*, a model research organism for amphibious fishes and evolution of hermaphroditism (Taylor 2012; Kelley et al. 2016), as sister to all other South American (Rivulidae) killifish species. New World (Rivulidae) and Old World (Nothobranchiidae, Aplocheilidae) killifishes are sister clades.

We modeled the evolution of annualism on the Bayesian phylogeny using maximum likelihood inference and stochastic character mapping as well as most parsimonious ancestral state reconstruction. Our findings support the hypothesis that annualism originated independently, more than once in both Africa and South America (Figure 1). We reject the reciprocal monophyly of annual and non-annual species in both New World and Old-World killifishes. Aplocheiloid killifishes began to diverge around 64 million years ago (56-72 mya HPD) around the Cretaceous-Paleogene boundary, and since that time, annualism has evolved up to seven times on two different continents (Figure 1).

### Molecular evolution of protein coding genes

Next, we compared rates of protein evolution in orthologous genes between annual and non-annual killifishes, measur(= d*N*/d*S*), in 1,234 genes for which we had both annual and non-annual fishes present in the alignment. To do so, we used codeml in the PAML package (Yang 2007) to obtain independent estimates of ω for annual and non-annual species for each alignment. We obtained higher mean estimates for annual killifishes, ω_annual_ = 0.23, relative to non-annuals, ω_non-annual_ = 0.18, with 29 genes that had either ω_annual_ or ω_non-annual_ higher than an arbitrarily set cutoff value of 3. These differences were significant in a Kolmogorov-Smirnov test (D=0.10349, P = 4.515e-06). We detected an increase in the proportion of alignments where ω_annual_ > ω_non-annual_ as we increased our level of stringency. This proportion went from 57% (699/1,223) when we considered all alignments other than those with ω_annual_ or ω_non-annual_ higher than 3, to 76% (161/211) when we restricted our analyses to genes where likelihood ratio tests found that ω_annual_ ≠ ω_non-annual_ at the 5% level, and to 85% (78/92) if we restricted our attention to genes where we adjusted P to be lower than 0.05 after a false discovery rate of 10%. These results are robust to changes in the cutoff value of ω, or to small changes in the tree topology. In cases where gene trees differed significantly from our species tree in Figure 1 based on the AU test (Shimodaira 2002), we obtained estimates of ω using both the species tree and the gene tree and found very small differences between them.

To summarize, our analyses pointed to 211 genes that had significantly different ω between annual and non-annual lineages. Within this gene set, a subset of 161 genes with higher d*N*/d*S* ratios in annuals (ω_annual_ > ω_non-annual_, Table 1) are fast evolving genes (FEGs) that are more likely to be positively selected genes (PSGs) in the annual lineages. Specifically, we find 11 PSGs (Table 2) that have ω_annual_ > 1 (FDR <0.05) and show strong signatures of positive selection across all sites in the alignment. GO term analyses reveal that many developmental and metabolic genes are represented in this group and thus they are likely pertinent to the annual life history. Furthermore, our FEGs with higher ω_annual_ are enriched for metabolic functions (Table 3, FDR < 0.05). Specifically, we show that our annual FEGs are enriched for nuclear genes controlling important aspects of metabolism and cell respiration including electron transport within mitochondria (OXPHOS, Figure 3), translation, and protein and nitrogen compound metabolism. We project the location of the 211 genes with significantly different ω between life-histories onto the chromosomes of the outgroup medaka (Supplementary Figure 1) showing distribution of these genes across the genome.

**Table 2.**
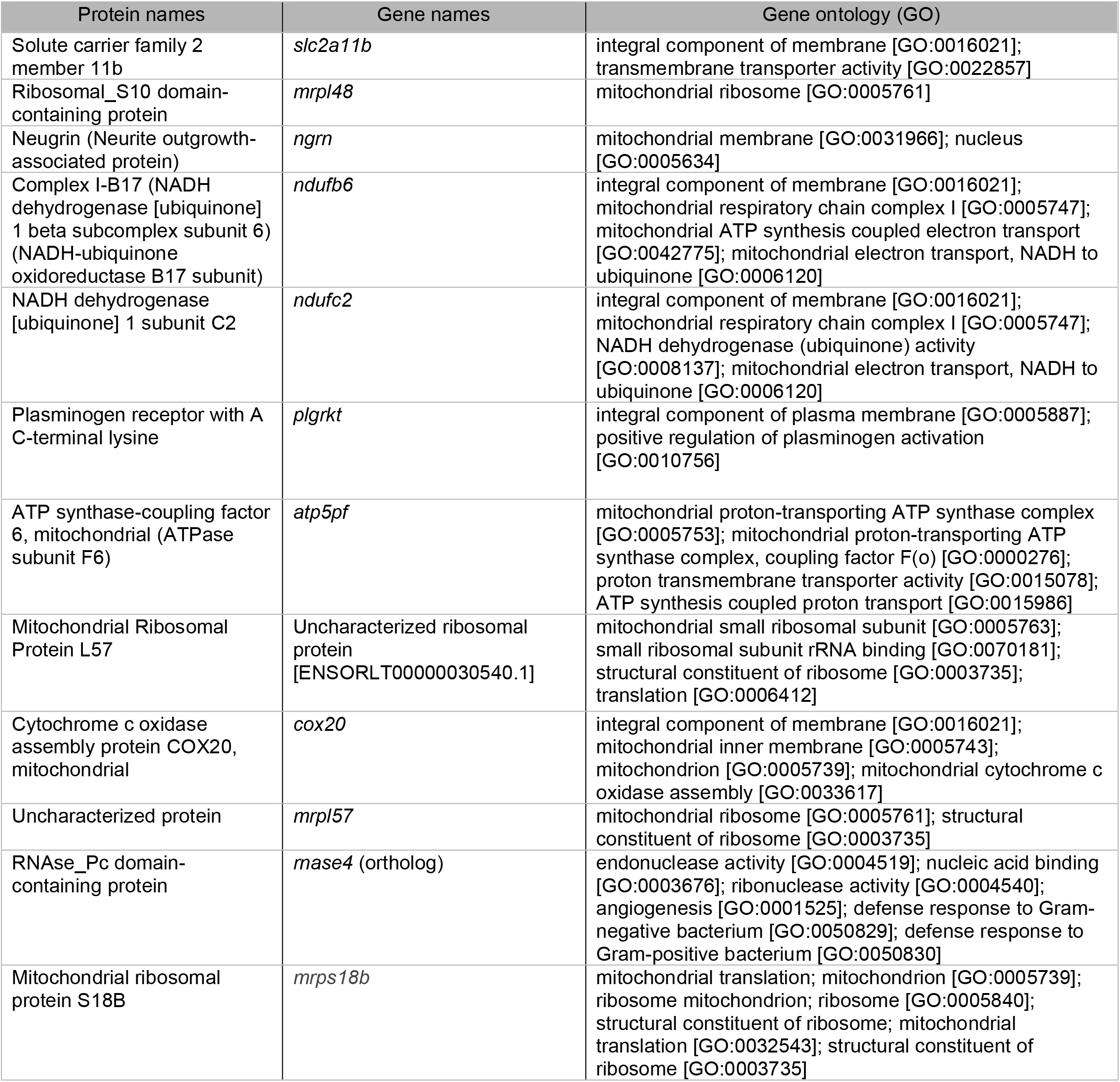
The 11 genes with the highest ω_annual_ with significantly higher d*N*/d*S* in annual lineages compared to non-annual lineages (ω_non-annual_). These FEGs are candidates for positive selection, and all have ω_annual_ > ω_background_; ω_annual_ > 1 (LRT >3.84). Gene ontology output from Uniprot.

| Protein names | Gene names | Gene ontology (GO) |
| --- | --- | --- |
| Solute carrier family 2 member 11b | *slc2a11b* | integral component of membrane [GO:0016021]; transmembrane transporter activity [GO:0022857] |
| Ribosomal_S10 domain-containing protein | *mrpl48* | mitochondrial ribosome [GO:0005761] |
| Neugrin (Neurite outgrowth-associated protein) | *ngrn* | mitochondrial membrane [GO:0031966]; nucleus [GO:0005634] |
| Complex I-B17 (NADH dehydrogenase [ubiquinone] 1 beta subcomplex subunit 6) (NADH-ubiquinone oxidoreductase B17 subunit) | *ndufb6* | integral component of membrane [GO:0016021]; mitochondrial respiratory chain complex I [GO:0005747]; mitochondrial ATP synthesis coupled electron transport [GO:0042775]; mitochondrial electron transport, NADH to ubiquinone [GO:0006120] |
| NADH dehydrogenase [ubiquinone] 1 subunit C2 | *ndufc2* | integral component of membrane [GO:0016021]; mitochondrial respiratory chain complex I [GO:0005747]; NADH dehydrogenase (ubiquinone) activity [GO:0008137]; mitochondrial electron transport, NADH to ubiquinone [GO:0006120] |
| Plasminogen receptor with A C-terminal lysine | *plgrkt* | integral component of plasma membrane [GO:0005887]; positive regulation of plasminogen activation [GO:0010756] |
| ATP synthase-coupling factor 6, mitochondrial (ATPase subunit F6) | *atp5pf* | mitochondrial proton-transporting ATP synthase complex [GO:0005753]; mitochondrial proton-transporting ATP synthase complex, coupling factor F(o) [GO:0000276]; proton transmembrane transporter activity [GO:0015078]; ATP synthesis coupled proton transport [GO:0015986] |
| Mitochondrial Ribosomal Protein L57 | Uncharacterized ribosomal protein [ENSORLT00000030540.1] | mitochondrial small ribosomal subunit [GO:0005763]; small ribosomal subunit rRNA binding [GO:0070181]; structural constituent of ribosome [GO:0003735]; translation [GO:0006412] |
| Cytochrome c oxidase assembly protein COX20, mitochondrial | *cox20* | integral component of membrane [GO:0016021]; mitochondrial inner membrane [GO:0005743]; mitochondrion [GO:0005739]; mitochondrial cytochrome c oxidase assembly [GO:0033617] |
| Uncharacterized protein | *mrpl57* | mitochondrial ribosome [GO:0005761]; structural constituent of ribosome [GO:0003735] |
| RNAse_Pc domain-containing protein | *rnase4* (ortholog) | endonuclease activity [GO:0004519]; nucleic acid binding [GO:0003676]; ribonuclease activity [GO:0004540]; angiogenesis [GO:0001525]; defense response to Gram-negative bacterium [GO:0050829]; defense response to Gram-positive bacterium [GO:0050830] |
| Mitochondrial ribosomal protein S18B | *mrps18b* | mitochondrial translation; mitochondrion [GO:0005739]; ribosome mitochondrion; ribosome [GO:0005840]; structural constituent of ribosome; mitochondrial translation [GO:0032543]; structural constituent of ribosome [GO:0003735] |

**Table 3.** Enriched biological functions of the 161 high ω_annual_ (FDR </=0.05) genes based on medaka gene annotations in PANTHER. See Supplementary Table 3 for full list.

| Biological Process | Fold Enrichment | FDR |
| --- | --- | --- |
| mitochondrial electron transport, NADH to ubiquinone (GO:0006120) | 41.45 | 3.07E-03 |
| mitochondrial cytochrome c oxidase assembly (GO:0033617) | 40.65 | 3.63E-02 |
| respiratory chain complex IV assembly (GO:0008535) | 37.75 | 4.19E-02 |
| aerobic electron transport chain (GO:0019646) | 24.66 | 6.96E-05 |
| mitochondrial ATP synthesis coupled electron transport (GO:0042775) | 24.18 | 6.30E-05 |
| oxidative phosphorylation (GO:0006119) | 22.73 | 2.51E-05 |
| ATP synthesis coupled electron transport (GO:0042773) | 22.02 | 8.10E-05 |
| electron transport chain (GO:0022900) | 19.79 | 2.03E-06 |
| respiratory electron transport chain (GO:0022904) | 17.87 | 2.34E-04 |
| apical junction assembly (GO:0043297) | 16.31 | 9.90E-03 |
| bicellular tight junction assembly (GO:0070830) | 16.31 | 9.38E-03 |
| tight junction organization (GO:0120193) | 15.19 | 1.23E-02 |
| tight junction assembly (GO:0120192) | 15.19 | 1.17E-02 |
| aerobic respiration (GO:0009060) | 15.15 | 1.16E-04 |
| ATP metabolic process (GO:0046034) | 13.76 | 1.79E-05 |
| cellular respiration (GO:0045333) | 12.93 | 2.90E-04 |
| energy derivation by oxidation of organic compounds (GO:0015980) | 10.14 | 1.18E-03 |
| generation of precursor metabolites and energy (GO:0006091) | 9.18 | 7.98E-05 |
| translation (GO:0006412) | 6.5 | 1.08E-03 |
| peptide biosynthetic process (GO:0043043) | 6.37 | 1.20E-03 |
| peptide metabolic process (GO:0006518) | 5.89 | 3.75E-04 |
| amide biosynthetic process (GO:0043604) | 5.2 | 6.39E-03 |
| cellular amide metabolic process (GO:0043603) | 4.29 | 7.01E-03 |

**Figure 3.**
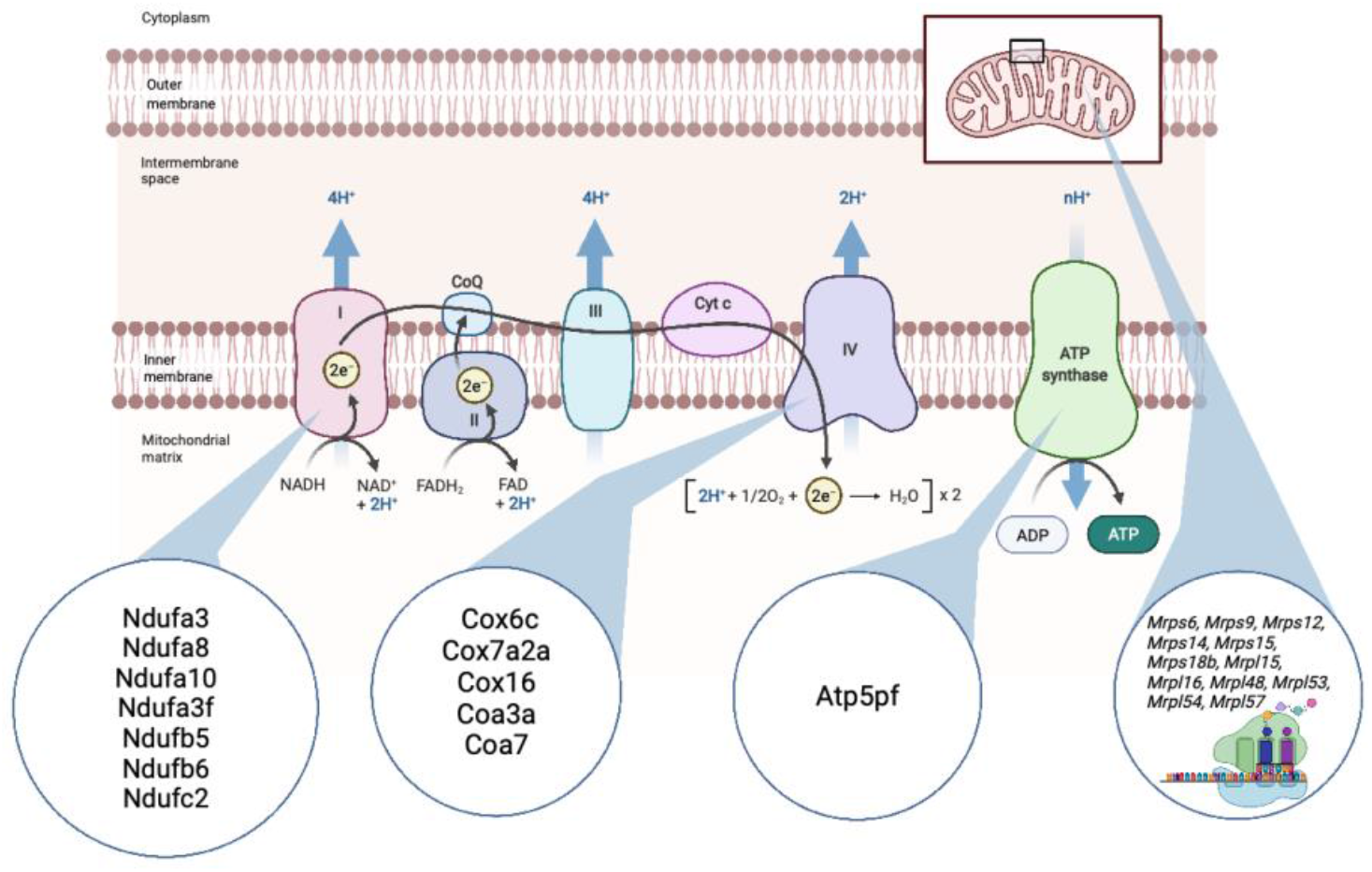
Higher ω_annual_ metabolic proteins. Many annual FEGs for OXPHOS proteins. Significantly higher ω_annual_ genes are part of complex I and IV of the electron transport chain as well as ATP synthase and mitochondrial ribosomal proteins. The oxidative phosphorylation pathway contains proteins such as Mrpl48, Ndufb6, Atp5pf, Ndufc2, Mrpl57, Cox20 with ω_annual_ > 1 across all sites indicating they have undergone strong positive selection. Adapted from “Electron Transport Chain”, by BioRender.com (2021). Retrieved from https://app.biorender.com/biorender-templates

### Identification of diapause-associated genes via codon and expression changes

To identify candidates undergoing both codon and regulatory changes involved in evolutionary transitions to extreme environments, we surveyed the literature to compare FEGs and PSGs found here to PSGs and differentially expressed genes (DEGs) associated with diapause in killifishes in other studies (Reichwald et al. 2015; Valenzano et al. 2015; Thompson and Ortí 2016; Wagner et al. 2018; Cui et al. 2019; Hu et al. 2020). We find 111 annual FEGs in our study that have not been previously identified as PSGs (Supplementary Table 4). In addition, we provide evidence to corroborate 20 genes with high ω_annual_ in our study that were previously proposed as PSGs by others (Supplementary Table 4). Additionally, we identify an overlap of 66 genes in our high ω_annual_ gene set with genes differentially expressed during DII and DIII that we term fast-evolving differentially expressed genes (FEDEGs, Supplementary Table 4,5; Figure 4). Aside from FEGs and PSGs, we identify 12 DEGs that consistently change expression in the same direction between DII and DIII and a non-dormant state in three different annual species (*Nematolebias whitei, Austrofundulus limnaeus, Nothobranchius furzeri*) representative of three different origins of annualism (Figures 1,4). These 12 DEGs include four genes (*ahnak, bahd1, csde1, and gtpbp4*) always upregulated during diapause, which we term core diapause DEGs or CDDEGs, and eight genes (*actl6a, adamts1, ccnb1, cdc20, ddx5, rcc1, ruvbl2, smarca5*) always downregulated in diapause (Figure 4, Supplementary Table 5).

**Figure 4.**
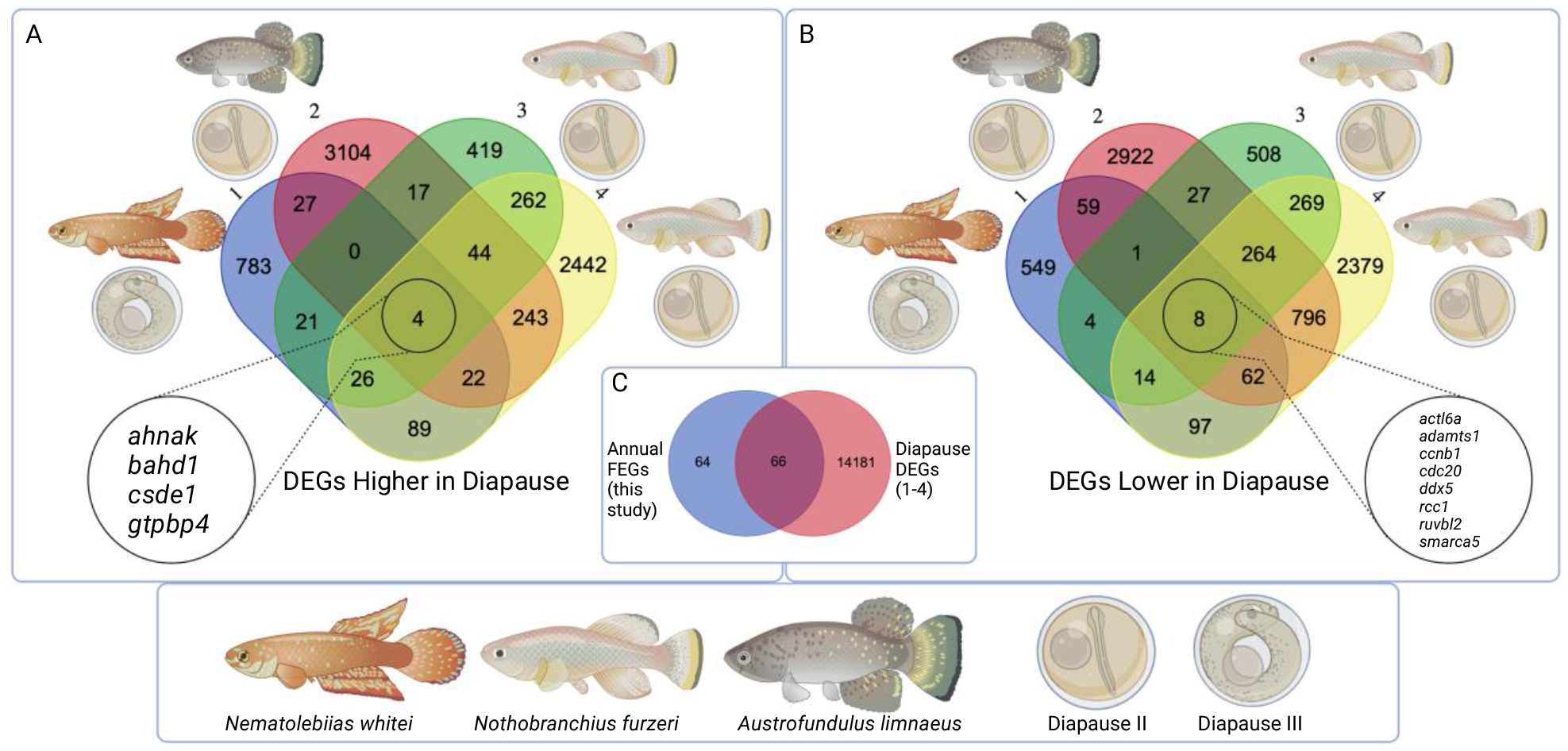
Diapause associated changes in gene expression across different studies of annual killifishes. Differentially expressed genes during 1.) DIII, Thompson and Ortí (2016); 2.) DII, Wagner et al. (2018); 3.) DII, Reichwald et al. (2015); and 4.) DII, Hu et al. (2020) on annual killifishes. These genes change expression in the same direction between diapause and equivalent non-dormant developmental stages (higher in diapause, A and lower in diapause, B) and 66 overlap with annual FEGs identified in our study (C). These studies include representative species from three origins of annualism: the Rio pearlfish, *Nematolebias whitei, Austrofundulus limnaeus*, and the turquoise killifish *Nothobranchius furzeri*. Created with BioRender.com.

## Discussion

Overall, our aplocheiloid killifish phylogeny represents the first genome wide inference to provide a robust, well-supported backbone for the evolutionary relationships and molecular evolution of this diverse model clade. This will allow future exploration of developmental and physiological novelties and potentially inform research on killifishes as emerging models for human disease (Cellerino et al. 2015; Harel, Benayoun, B. Machado, et al. 2015; Reichwald et al. 2015; Valenzano et al. 2015; Thompson and Ortí 2016). Phylogenetic hypotheses for this group are relatively stable with no evidence of incomplete lineage sorting, with support for multiple origins of annualism in both Africa and South America. Interestingly, we also find that the earliest likely divergence of the 3 killifish subclades (Figure 1) was ∼72 mya and around this time, the continental regions of South America, Africa, Madagascar, and India were already well separated. Since vicariance would be better supported with earlier divergence times, the current, global distribution of these fishes is best explained by dispersal events. Although all extant species of Aplocheiloid killifishes inhabit freshwaters, there is strong evidence for intercontinental and marine dispersal in this group of fishes (Ponce de Leon et al. 2014; Cui et al. 2021). We focus our analysis here on the molecular evolutionary underpinnings of up to seven convergent origins of annualism in killifishes. However, we also note that our phylogenomic analysis also reveals other types of convergence in this clade. For instance, our phylogeny rejects the monophyly of non-annual killifishes in Rivulidae that were previously classified in the genus *Rivulus* and hypothesized by others to be monophyletic (Murphy and Collier 1996; Huber 2012). The relationships within our phylogeny support a previously proposed taxonomic break-up and renaming of the genus (Costa 2011). These species are slender bodied, often amphibious, air-breathing fishes showing sexual dichromatism with females possessing a dark pigmented spot on the caudal peduncle near the tail. Consequently, our results illustrate that this “*Rivulus*” phenotype is another example of convergent evolution or even retention of ancestral characters. Thus, our phylogeny will serve as a foundation for future comparative work on the molecular evolutionary basis of other morphological and physiological adaptations besides annualism in this important model clade.

Annual killifishes must undergo drastic metabolic changes and challenges during their development including exposure to low oxygen conditions (Podrabsky et al. 1997). Metabolism is greatly depressed during diapause (Podrabsky and Hand 1999) and anoxia tolerance has evolved in conjunction with developmental delay with hypoxia initiating DI (Peters 1963) and anoxia tolerance during DII (Podrabsky et al. 2007). Low oxygen conditions are also hypothesized to trigger hatching and termination of DIII during habitat inundation in seasonal environments (Wourms 1972c; Dimichele and Taylor 1980; Dimichele and Taylor 1981; Dimichele and Powers 1984; Podrabsky and Hand 1999). Thus, it is expected that OXPHOS genes will evolve under selection pressures in fishes with varying degrees of aerobic or anaerobic challenges (Zhang and Broughton 2015). We find fast evolving genes encoding OXPHOS proteins are implicated in the transition to a seasonal life history strategy. We show that these OXPHOS and mitochondrial protein synthesis genes with critical mitochondrial function (Figure 3) have much higher ω_annual_ than the rest of the annual FEGs (Table 2).

The same electron transport protein complexes have been convergently modified in other metabolically-challenged vertebrates living in extreme, low oxygen environments. For example, subterranean, marine, and high elevation mammals (Tian et al. 2017) as well as high-elevation hummingbirds (Lim et al. 2019) and loaches (Wang et al. 2016) have higher ω and/or positive selection acting on OXPHOS proteins associated with NADH ubiquinone oxidoreductase (complex I) and ATPase (complex IV) (Figure 3). Interestingly, the opposite trend is observed in fishes that live in fast water flow with high dissolved oxygen concentrations (Lu et al. 2019). These fish experience a relaxation of purifying selection on mitochondrial OXPHOS genes. All this evidence, along with significant enrichment (Table 3) of high ω_annual_ genes involved in OXPHOS as well as four OSPHOS genes with ω_annual_ >1 (Table 2) further suggests the importance of these genes for the convergent change of physiology in extreme environments where oxygen is limiting, and hypoxia/anoxia tolerance is necessary for survival. More specifically, it illustrates specific pathways that are more likely to change in response to extreme environments as shown by the repetitive, convergent increases in dN/dS across divergent vertebrate lineages and suggests a core set of genes that convergently and rapidly evolve in harsh conditions.

We also find many genes that control developmental phenotypes include critical transcription factors, enzymes, and signaling proteins such as *gchfr, hoxd4a, foxc1, tbx16, prrx1, rax, her4*.*2, hand2, bhlha9*, undergo rapid evolution in annual species. None of these genes have been previously associated with positive selection in annual killifishes. These genes include specific developmental genes which evolve in correlation to the annual or seasonal phenotype and diapause. Specifically, *forkhead box c1* (*foxc1)* and *T-box transcription factor 16* (*tbx16*) genes have developmental roles such as regulation of somitogenesis (Topczewska et al. 2001; Morrow et al. 2017). Diapause II occurs during somitogenesis, and these genes may have important functions during DII present in all annual lineages. GTP cyclohydrolase 1 feedback regulator (*gchfr)* is a FEDEG, significantly changing in expression during killifish diapause (Supplementary. Table 5; (Reichwald et al. 2015; Wagner et al. 2018) and may be associated with both metabolic and developmental changes. Interestingly, *gchfr* is involved in negative regulation of GTP cyclohydrolase, an enzyme which, when mutated, can lead to developmental delay in humans (Longo 2009). Furthermore, *gchfr* is also a target of the AhR pathway which is upregulated and results in lethal developmental anomalies in Atlantic killifish (*Fundulus heteroclitus*) exposed to pollutants (Dubansky et al. 2013). Consequently, we hypothesize *gchfr* may have undergone both regulatory expression evolution and protein structure evolution that affect developmental pathways and developmental delay in annual killifishes with multiple implications for adaptation to extreme environments. Further study is needed to elucidate the function of these developmental genes in diapausing embryos, but our results highlight the utility of annual killifishes for exploring the relationship between metabolism and development.

Our analyses identify a total of 111 genes with high ω_annual_ (Supplementary Table 4), significantly more than previous studies, likely due to the large number of species compared. Taxon sampling can change inferences regarding molecular evolution (Sahm et al. 2016) as these genes were also included in previous studies but not undergoing positive selection in annual species. Previous studies focused on PSGs or mutations in one or two annual lineages (Sahm et al. 2016). For example, it has been reported that African annual killifish (e.g. *Nothobranchius* and *Callopanchax* species) have high rates of mtDNA mutation and relaxed selection (Cui et al. 2019). Additionally, others reported evidence for positive selection of mitochondrial-associated nuclear genes in the American annual species *Austrofundulus limnaeus* (Wagner et al. 2018). Our results illustrate the repetitive faster protein evolution correlated with specific life histories by using a branch model to look at evolutionary rate across many lineages. Our findings point to fast evolution of metabolic genes across many species with similar life histories and show this evolutionary pattern is associated with dormancy and rapid maturation and growth. Besides changes in protein codon evolution, our meta-analysis reveals 12 DEGs that consistently change expression in the same direction during DII and DIII (Figure 4). We find four genes (CDDEGs) always upregulated during diapause and 8 genes always down regulated in diapause (Supplementary Table 5). This pattern is consistent even though it is observed in two different diapause stages (DII and DIII) and in three different species each of which independently evolved an annual life history. DII and DIII occur at drastically different developmental time points when the embryo has a completely different set of anatomical features (Wourms 1972a). However, since specific phenotypes are required for diapause such as metabolic depression and cessation of cell cycle or proliferation, some of the same key genes may be repeatedly activated or repressed to maintain dormancy (Thompson and Ortí 2016). Killifish diapause is an active state where genes are still expressed and epigenomic and gene expression changes still occur (Thompson and Ortí 2016; Hu et al. 2020). Therefore, genes being upregulated during diapause are expected to have functions pertaining to maintaining a dormant state. Some of these genes may be activated to repress normal functions that create the procession of development.

Within our CDDEGs, we find consistent changes in expression of genes related to cell cycle and cell proliferation. For example, the neuroblast differentiation-associated protein Ahnak is upregulated in DII in the turquoise killifish, *Austrofundulus*, and DIII in the Rio Pearlfish. The *ahnak* gene functions as a tumor suppressor (Lee et al. 2014) and may reduce cell proliferation during DII and DIII. Similarly, downregulated genes include *cyclinB1* (*ccnb1*), *cell division cycle protein 20* (*cdc20*), and *regulation of chromatin condensation 1* (*rcc1*), all of which have functions in mitotic progression and cell proliferation and when overexpressed increase probability of tumorigenesis (Cekan et al. 2016; Otto and Sicinski 2017; Ren et al. 2020). This further highlights the potential for research on gene regulatory networks involved in annual killifish adaptations to inform research on pathways involved in human pathologies. The four CDDEGs do not show evidence of positive selection, rapid evolution, or slower evolution in annual lineages suggesting that diapause phenotype is modulated by both changes in coding sequences as well as transcript expression levels that could be subject to differential regulation in annual species.

### Conclusions

While many previous studies have focused on a single species or subclade of killifishes, our analyses span the entire diversity of aplocheiloids and consequently uncover more candidate loci potentially involved in the adaptation to extreme environments via changes in both protein sequence evolution and gene expression. Studies such as ours are important in the face of global change and can highlight genes and pathways that are the focus of selection and convergent adaptation to extreme environments and provide candidates for exploration with functional genomic tools. Overall, our results suggest that annual killifishes evolve faster than their non-annual counterparts and that shifts in rates of molecular evolution are likely convergent and adaptive for ephemeral habitats. Annual killifishes have more rapidly evolving developmental and OXPHOS proteins likely due to the unique ontogenetic and physiological demands imposed by diapause and anoxia exposure. Our study points to proteins and pathways that are convergently evolving in highly disparate vertebrate groups living in harsh conditions. Stephan J. Gould’s contingency theory proposed that if we “replay the tape” of life, evolution could be highly contingent and unpredictable (Gould 1989). Based on our findings here, we rather reject this hypothesis and support evolutionary determinism at a global scale in annual killifishes, showing that annual killifishes and other extremophiles “replay” their adaptation to similarly harsh environments using similar molecular evolutionary signatures.

## Materials and Methods

### Sample preparation

RNA was extracted from frozen, RNAlater preserved whole fish or embryos (Supplementary Table 1) with Trizol or a Qiagen RNeasy plus extraction kit according to manufacturer’s instructions. RNA was stabilized with GenTegra-RNA tubes following manufacturer’s instructions and sent to the Beijing Genome Institute (BGI) for RNA-Seq. Samples were prepared with a TruSeq^®^ RNA Sample Preparation Kit (Illumina, San Diego, CA, USA) to construct Illumina cDNA libraries according to the manufacturer’s protocol. About 3-Gb paired-end raw reads with a length of 100bp were generated for each sample using Illumina HiSeq 4000 platforms (Illumina, San Diego, CA, USA) at BGI-Tech (BGI, Shenzhen, Guangdong, China).

### Sequencing, quality control, and assembly

Raw sequence data was demultiplexed and quality controlled via Trimmomatic v 0.32 (Bolger et al. 2014) with the following parameters: PE, ILLUMINACLIP:2:30:7, HEADCROP:13, SLIDINGWINDOW:7:15, MINLEN:25. A transcriptome was assembled de novo for each species with Trinity 2.1.1 (Grabherr et al. 2011; Haas et al. 2013) with the following parameters: --seqType fq --normalize_reads. The resulting transcriptome was filtered for ribosomal RNA and microbial contaminants using SortMeRNA v 2.0 (Kopylova et al. 2012) against a non-redundant database compiled from exons and mRNA of the annual killifishes *A. limnaeus* and *N. furzeri* genomes and transcriptomes available in Genbank. Transdecoder (Grabherr et al. 2011; Haas et al. 2013) was used to find open reading frames in each transcriptome. For species that were already sequenced, CDS data was extracted from available transcriptomes (as in *K. marmoratus* with the described Trinity pipeline and Transdecoder) or from available genomes with cufflinks-2.2.1 (Trapnell et al. 2012) gffread feature.

The PhyloUtensils (https://github.com/ballesterus/Utensils) and UPhO (Ballesteros and Hormiga 2016) software packages were used to analyze data and identify orthologs. Putative homologs between and within species transcriptomes were identified with an initial all vs all BLASTP with an e-value cutoff of e-3 and then clustering with MCL (Ballesteros and Hormiga 2016) using only hits with an e-value of e-10. Orthogroup alignments were made with results from homologs with 5+ taxa at I =1.4 with TranslatorX (Abascal et al. 2010) using the following parameters: -c 1 -t T -p F. TranslatorX found the best open reading frame, and these alignments were input to pamatrix.py with default settings for al2phyo.py. RAxML v. 8.1.23 (Abascal et al. 2010) was used to create trees for each orthogroup alignment partitioned by codon with the following parameters in parallel, partitioned by codon to get orthogroup trees: parallel -q -s -m GTRCAT -f a -p 767 -x 97897 -#100 -n.

We used UPho.py to extract orthologs from each orthogroup tree with a minimum of 7 taxa and with clades of 75%+ bootstrap support. Get_Fasta_from_Ref.py (https://github.com/ballesterus/UPhO) was used to pull out nucleotide fasta sequences for 2557 orthologs from the Transdecoder output. Isoforms were not chosen for analysis (i.e. the longest one). This is because it is difficult to tell apart duplicates, alleles, and isoforms in Trinity output without reference genomes for each species. Using the UPhO pipeline, they behave as in-paralogs which were not kept in our analysis (Ballesteros and Hormiga 2016). These sequences were realigned with only their orthologous sequences from other species via TranslatorX using the same parameters as before. Then, we ran PaMATRIX.py and aplo2phy as follows with -t = 7. This resulted in 2557 alignments. Only alignments containing aplocheiloid killifish species were kept. These were edited manually, removing any spurious regions or sequences of homology. Alignments with at least 5 taxa after manual check and at least one aplocheiloid were kept. This resulted in 1,302 alignments. A BLASTN search confirmed they had CDS homologs in the medaka Ensembl genome assembly (Supplementary Table 2). These 1302 single copy, orthologous gene loci alignments were concatenated into a super matrix totaling 852,927 bp used for phylogenetic analyses.

### Phylogeny construction and character analyses

Using our super matrix of 1302 core orthologs, we inferred a time calibrated phylogeny using BEAST 2.5.3 (Bouckaert et al. 2019) linking clock models and trees with an MCMC chain length of 785,344,000 sampling every 1000 states for a total of 785,344 samples. Data was partitioned by codon and a GTR site model with four rate categories was used. We used a birth-death tree prior and a relaxed lognormal clock with three clock calibrations at three different nodes. Since fossils are extremely rare for these small freshwater fishes (Altner and Reichenbacher 2015), we used estimated time values from timetree.org (Kumar et al. 2017) at the following nodes: MRCA killifishes and *Danio* (230 mya, offset = 223), MRCA killifishes and *Oryzias* (93 mya, offset =86), and MRCA killifishes and Poecilia (75 mya, offset = 68). Upon completion of the MCMC chain, tree annotator was used to obtain the maximum clade credibility tree with 30% of trees discarded as burn-in. The same ortholog data matrix used in the Bayesian analysis was also input into RAxMLHPC8 with the following parameters -T 4 -f a -N autoMRE -m GTRGAMMA -p 12345 -x 12345 -o Danio_rerio to infer a ML species tree.

The time-calibrated tree served as an input to estimate the evolution of annualism via stochastic character mapping. We created a life history character matrix where annualism was coded as a binary character due to presence or absence of DII as described in (Furness et al. 2015). Annualism transition rates were modeled with the “All Rates Different” (ARD) model in the phytools R package (Revell 2012) that allows forward and reverse rates of character evolution to differ. We then used the phytools simmap function to infer and plot the posterior probability of evolving an annual life history across the branches of the phylogeny with 100,000 simulations (Figure 1). Additionally, our life history character matrix was used to infer a most parsimonious ancestral state reconstruction of this trait on the phylogeny. This result allowed us to characterize annual lineages based on the number of origins of annualism and this was used as an input to characterize_lineage.py to parse annual and non-annual lineages as foreground and background branches respectively for subsequent d*N*/d*S* analyses (described below). A total of 1,234 alignments with at least 1 annual species (out of 1302) were used for modeling rates of protein evolution.

To test the robustness of our phylogeny, we also conducted a species tree analysis to rule out confounding effects of incomplete lineage sorting. We inferred phylogenetic relationships of our 1,302 core killifish alignments with IQTree-1.6.1 (Nguyen et al. 2015) using maximum likelihood with the model of nucleotide substitution selected with the standard model selection from IQ-Tree to obtain gene trees for each ortholog. The gene trees were analyzed with ASTRAL II (Mirarab and Warnow 2015) to obtain a species tree which eliminated any confounding effects of incomplete lineage sorting on the inference of the species tree.

### Rate of evolution and gene function

Our 1,302 loci were confirmed to be protein coding genes via BLASTn against the medaka coding sequences in Ensembl 94 (Supplementary table 2). For each alignment of orthologous sequences, we explored variation in the rate of molecular evolution between annual and non-annual lineages using the codon-based maximum likelihood framework introduced by Goldman and Yang (1994) as implemented in the program codeml from the PAML 4.9 (Goldman and Yang 1994) package. Briefly, for a given tree and alignment, we compared the ratio of the rate of non-synonymous substitutions per non-synonymous site (dN) and the rate of synonymous substitutions per synonymous site (dS). For each alignment of orthologous sequences, we compared a null model where all branches of the tree were constrained to have the same ω (=d*N*/d*S*) with a model where branches were assigned to two different groups, annual and non-annual, each with an independent estimate of ω, ω_annual_ and ω_non-annual_. Because the null model is nested within the alternative model, we checked whether the estimated differences in ω were significant with a likelihood ratio test with critical values derived from a chi-square distribution with 1 degree of freedom, as suggested by the PAML manual (http://abacus.gene.ucl.ac.uk/software/pamlDOC.pdf, last accessed in March 2019).

To explore the potential role of tree misspecification in the analyses, for each candidate gene alignment of orthologous sequences, used the topology obtained from IQTree (Nguyen et al. 2015), and tested whether the resulting tree topology diverged from the expected organismal phylogeny using the approximately unbiased test (Shimodaira 2002) with 10000 RELL replicates as implemented in IQTree-1.6.1. In cases where the two trees differed significantly, we ran the analyses with each of the two trees. We used a model selected by tree reconstruction, 1000 replicates of ultrafast bootstrap (Nguyen et al. 2015), and 1000 replicates in the Shimodaira-Hasegawa approximate likelihood-ratio test and approximate Bayes test (Thi Hoang et al. 2017). A concatenated phylogeny including all species studied (Figure 1) was pruned of any species not in the alignment of interest, and the resulting tree was stripped of branch lengths and annual labels applied to appropriate branches and species. Topology tests were run in IQTree-1.6.1, comparing the maximum likelihood tree to the pruned tree with the same model used in tree construction, 10000 RELL replicates in the approximately unbiased test (Anisimova et al. 2011) and weighted-KH and weighted-SH tests. The subsequent annual-labeled best tree was then used in codeml, running branch analysis with Seqtype=1, model=2, ncatG=8, RateAncestor=1, and remaining parameters at default settings; and s sites analysis was run on the unlabeled version, with Seqtype=1, model=0, NSsites=0 1 2 3, ncatG=8, RateAncestor=1, and remaining parameters at default settings.

We then compared the ω distributions between these two sets of species using a Kolmogorov–Smirnov test. Because short branches can lead to unreliable estimates of ω, we either 1-removed ω estimates higher than 3 from the analyses, or 2-arbitrarily changed ω higher than 3 to 3. These tests were robust to changes in the arbitrary value of ω set as boundary (2, 5, or 10).

Gene alignments that had significant differences in d*N*/d*S* between annuals and non-annuals were collapsed based on a 0% majority consensus where the most common bases and fewest ambiguities were kept. These consensus sequences were searched against the medaka Ensembl Protein database 94 with BLASTX to find the best scoring hit. Gene ontology terms were mapped to these ensemble protein identities for medaka with the PANTHER (Mi et al. 2016) online classification system available through the Gene Ontology Consortium. A statistical overrepresentation test using GOs mapped to medaka was conducted with a false discovery rate (FDR) cutoff of 0.05 on the complete GO biological processes database via PANTHER for genes that had faster and slower rates of evolution in annual lineages.

### Comparison of fast evolving, positively selected, and differentially expressed genes

We also conducted a meta-analysis to examine overlap in genes that had similar differential expression patterns in diapause and higher ω_annual_ across this and other studies (Mi et al. 2016). We summarized gene overlap between studies by classifying species that were described as annual (Reichwald et al. 2015; Valenzano et al. 2015; Thompson and Ortí 2016; Sahm et al. 2017; Wagner et al. 2018; Cui et al. 2019; Hu et al. 2020) or “semi-annual” (Cui et al. 2019) as annuals and other species as non-annuals. Annotated gene symbols were mined from each study and DEGs (FDR < 0.05) from Thompson and Ortí (2016) were reannotated with best BLASTX hit from the medaka Ensembl Protein database 94. Common gene overlap of characterized genes is summarized in supp. tables 4 and 5.

## Acknowledgements and Funding Information

We thank J. Ballesteros, L. Hughes and everyone in the Ortí, Braasch, and Podrabsky labs for their feedback on analysis and the manuscript. We thank A. Terceira for kindly providing the fish photos and C. Olivera, J. Henriques, and R. Britzke for help with tissue collection. This work was supported by by the NSF BEACON Center for the Study of Evolution in Action (Cooperative Agreement No. DBI-0939454), the National Science Foundation (DEB-1406537), the National Geographic Society, the Cosmos Club of Washington D.C., and the American Killifish Association.

